# Genomic and functional analysis of *rmp* locus variants in *Klebsiella pneumoniae*

**DOI:** 10.1101/2024.05.28.596137

**Authors:** Margaret M.C. Lam, Stephen M. Salisbury, Logan P. Treat, Ryan R. Wick, Louise M. Judd, Kelly L. Wyres, Sylvain Brisse, Kimberly A. Walker, Virginia L. Miller, Kathryn E. Holt

## Abstract

**Background:** *Klebsiella pneumoniae* is an opportunistic pathogen and a leading cause of healthcare-associated infections in hospitals, which are frequently antimicrobial resistant (AMR). Exacerbating the public health threat posed by *K. pneumoniae*, some strains also harbor additional hypervirulence determinants typically acquired via mobile genetic elements such as the well-characterised large virulence plasmid KpVP-1. The *rmpADC* locus is considered a key virulence feature of *K. pneumoniae* and is associated with upregulated capsule expression and the hypermucoid phenotype, which can enhance virulence by contributing to serum resistance. Typically such strains have been susceptible to all antimicrobials besides ampicillin, however the recent emergence of AMR hypermucoid strains is concerning.

**Methods:** Here, we investigate the genetic diversity, evolution, mobilisation and prevalence of *rmpADC*, in a dataset of 14000 genomes from isolates of the *Klebsiella pneumoniae* species complex, and describe the RmST virulence typing scheme for tracking *rmpADC* variants for the purposes of genomic surveillance. Additionally, we examine the functionality of representatives for variants of *rmpADC* introduced into a mutant strain lacking its native *rmpADC* locus.

**Results:** The *rmpADC* locus was detected in 7% of the dataset, mostly from genomes of *K. pneumoniae* and a very small number of *K. variicola* and *K. quasipneumoniae*. Sequence variants of *rmpADC* grouped into five distinct lineages (*rmp1, rmp2, rmp2A, rmp3* and *rmp4*) that corresponded to unique mobile elements, and were differentially distributed across different populations (i.e. clonal groups) of *K. pneumoniae*. All variants were demonstrated to produce enhanced capsule production and hypermucoviscosity.

**Conclusion:** These results provide an overview of the diversity and evolution of a prominent *K. pneumoniae* virulence factor and support the idea that screening for *rmpADC* in *K. pneumoniae* isolates and genomes is valuable to monitor the emergence and spread of hypermucoid *K. pneumoniae*, including AMR strains.

## Background

A distinct pathotype of *Klebsiella pneumoniae*, often referred to as hypervirulent *K. pneumoniae* (hvKp), poses a significant public health challenge outside of clinical settings where it causes severe and sometimes life-threatening infections^1–3^. These are regarded as being distinct from ‘classical *Kp*’ strains, which typically cause opportunistic infections mostly within healthcare settings, and are often multidrug-resistant (MDR). Community-acquired infections can arise in otherwise healthy and immunocompetent individuals, although there are reportedly associations with comorbidities such as diabetes^4^. Common examples of infections include pyogenic liver abscess, endophthalmitis, pneumonia and meningitis, but they can also present as metastatic, multisite infections. Earlier reports of hypervirulent, community-acquired infections were largely confined to countries in Eastern Asia. Cases are now being more widely reported in other regions including Europe, North America and Australia, although often associated with individuals of East Asian descent^5^.

The overwhelming majority of hvKp infections are associated with strains from distinct genetic backgrounds or lineages; these include clonal groups CG23, CG25, CG65, CG66, CG86 and CG380^6,7^. Several features are considered hallmark characteristics of these hvKp strains^7,8^. Most produce a K1 or K2 capsule, encoded by the *cps* (K) loci KL1 and KL2, respectively, and O1 lipopolysaccharide (OL1 locus). Many hvKp strains also exhibit hypermucoviscosity (HMV), which is defined by a positive string test and/or low OD_600_ measurements in a sedimentation assay, and is associated with presence of the *rmpADC* locus and/or *rmpA2* gene. Lastly, most hvKp also synthesise the siderophores aerobactin (*iuc* locus) and salmochelin (*iro* locus), in addition to the intrinsic siderophore enterobactin (*ent* locus)^7,8^. The acquired siderophore loci, *rmpADC* and *rmpA2* (which also appears to be part of a locus including homologs of *rmpD* and *rmpC*), are typically mobilised by the large *K. pneumoniae* virulence plasmids (KpVP) that have been stably maintained for over 100 years in some HvKp clones including CG23 and CG86, although *iro* and *rmpADC* can also be mobilised via the chromosomal integrative conjugative element ICE*Kp1*^9–11^.

In *K. pneumoniae,* the hypermucoid or HMV phenotype has long been associated with the gene *rmpA* although the exact mechanisms leading to the phenotype were unknown^12–14^. Based on observations and experimental evidence from earlier studies using knockout mutants, it was proposed that the expression of *rmpA* upregulated expression of *cps* thereby increasing capsule production and subsequently resulting in HMV. Subsequent work has confirmed that the *rmpA* LuxR-like transcriptional regulator is part of a larger operon (herein called *rmp*) together with the *rmpD* and *rmpC* genes located downstream^15,16^. This work has further clarified that enhanced capsule expression and HMV are two discrete features requiring *rmpC* (regulator of *cps*) and *rmpD* (encoding a small protein required for HMV), respectively. While HMV can be attained in the absence of elevated *cps* expression (i.e. in a *rmpC* mutant), it was not observed in capsule-defect mutants, suggesting that HMV does rely on the presence of some capsular components but these need not be hyperexpressed. Accordingly, it was recently shown that HMV is driven through elongation of the capsule polysaccharide chain, driven by a direct interaction between RmpD and Wzc (transmembrane protein), and presumptive indirect interaction with Wzy (capsule repeat unit polymerase), both of which are components of the core capsule synthesis and export machinery^17,18^. The *rmpA* and *rmpA2* genes appear to be frequently subjected to insertions or deletions (indels) within a poly(G) tract that consequently encode a truncated and presumably non-functional product, and this has been suggested as a mechanism by which differential expression of the two genes is achieved^14^.

The presence or absence of *rmpA* in clinical isolates has been investigated in dozens of studies focused on hypervirulent infections, and *rmpA* has been identified as one of several biomarkers for distinguishing hvKp strains from non-hvKp^19^. However, their detection shows a variable degree of correlation with HMV as measured by string test (51-98%). The functional impact of allelic variation in the *rmp* locus genes has not yet been explored, although this is likely to be important for explaining or predicting HMV based on *rmp* locus sequences. Here we investigate the genetic diversity and distribution of the *rmp* locus in the *K. pneumoniae* species complex (KpSC), identify key variant lineages of *rmp* and their associated mobile genetic elements (MGEs), and demonstrate that representatives of each of the lineages are able to induce HMV and elevated capsule production when introduced into an hvKp isolate lacking its native *rmp* locus (KPPR1S *rmp*).

## Methods

### Genome sequences and genotyping

The initial screening for *rmpADC* was conducted on the same 2733 *K. pneumoniae* species complex (KpSC) genomes included in a 2018 study^10^ examining the aerobactin- and salmochelin-encoding loci *iuc* and *iro*, which are often co-localised with *rmp* on the same mobile genetic elements. The SRST2-table-from-assemblies.py Python script (github.com/rrwick/SRST2-table-from-assemblies) was used to screen these assemblies for the presence of existing *rmpA* alleles from the virulence database on BIGSdb-*Kp* (37 alleles as of October 2020; http://bigsdb.pasteur.fr/klebsiella/klebsiella.html) along with the reference *rmpD* and *rmpC* sequences from Walker *et al.*^15,16^ with BLAST+ v2.2.21, and novel alleles extracted with the --report_new_consensus flag. Unique alleles from 160/2733 genomes with an intact *rmp* locus (defined as those in which all genes in the locus could be assigned an allele) were assigned allele numbers, and unique allele combinations were used to define unique ‘*rmp* sequence types (RmSTs)’ using a multi-locus sequence typing (MLST) approach. These *rmpADC* alleles and RmST profiles were then incorporated along with *rmpA2* alleles into the genotyping pipeline Kleborate v2.0.0, and applied to screen 13,156 publicly available *Klebsiella* genomes^20^. From this dataset of 13,156 genomes, 944 genomes positive for *rmp* and/or *rmpA2* (including genomes with an incomplete *rmp* locus) were included for analysis in this study (see **Additional File 1** for genome accessions, isolate metadata and genotyping information). We also included *rmp*-positive genomes from three additional datasets: 4/208 isolates collected at an Australian hospital in 2002^21^, in addition to 36/392 isolates from the same hospital in 2020, and 8/276 isolates from the Burden of Antibiotic Resistance in Neonates from Developing Societies (BARNARDS) network^22^. Novel *rmpADC* alleles and RmST profiles were added to the BIGSdb-*Kp* database. Clonal groups, and designations of clones as hypervirulent or MDR, were done using previously defined ST-CG assignments^23^.

### Phylogenetic analyses

For each unique RmST, an alignment of the concatenated *rmpA*, *rmpD* and *rmpC* sequences was generated with MUSCLE v3.8.31, and used as an input for maximum likelihood phylogenetic inference with RAxML v8.2.9^24^ run five times with the generalised time-reversible GTR+gamma model. Similarly, phylogenies were also generated for *rmpA, rmpD* and *rmpC* genes individually using the same approach described for RmST. *Rmp* lineages were defined based on monophyletic groups of RmSTs mobilised by a unique mobile element (see below).

Phylogenetic trees for genomes belonging to hypervirulent clonal groups CG23, CG65 and CG86 were generated using cgMLST implemented in Pathogenwatch (pathogen.watch)^25^.

### Comparison of *rmp* genetic contexts

Contigs containing *rmp* were manually inspected in Bandage v0.8.1^26^ to determine whether the locus was located on the chromosome or on a previously described virulence plasmid (KpVP-1 reference pSGH10 with accession CP025081.1, or KpVP-2 reference Kp52.145 plasmid II with accession FO834905.1). For any assemblies where the *rmp*-containing contig did not match to any of the contexts described above, the contig sequence was screened against the NCBI non-redundant nucleotide database and top hits noted. A single annotated representative (complete sequence where possible) was selected for each *rmp* lineage to compare the overall genetic structures of the neighbouring regions of *rmp* (i.e. up to 15 kbp upstream and downstream of *rmp*). Annotations were performed with Prokka v1.14.6^27^ and comparisons between the annotated genes were visualised with clinker v0.0.21 (github.com/gamcil/clinker)^28^.

### Bacterial strains, plasmids and growth conditions

The *K. pneumoniae* strains and plasmids used in this study are listed in **Table 1**. The strains from which representative *rmp* was amplified were as follows: SGH10 for *rmp1*, 52.145 for *rmp2*, NCTC 13669 for *rmp2A*, KPPR1S for *rmp3*, and NCTC 1936 for *rmp4*. Strains were grown at 37°C in LB medium (10 g tryptone, 5 g yeast extract, 10 g NaCl). Saturated overnight cultures were diluted to an OD_600_ of 0.2 and grown for 5.5 hours. Antibiotics were used where appropriate: kanamycin (Kan), 50 μg/ml; rifampin (Rif), 30 μg/ml; spectinomycin (Sp), 50 μg/ml. All plasmids were introduced into *K. pneumoniae* by electroporation as previously described^15^. For expression of genes cloned into the pMWO-078 vector, 100 ng/ml anhydrous tetracycline (aTc) was added to the medium at the time of subculture.

**Table 1.**
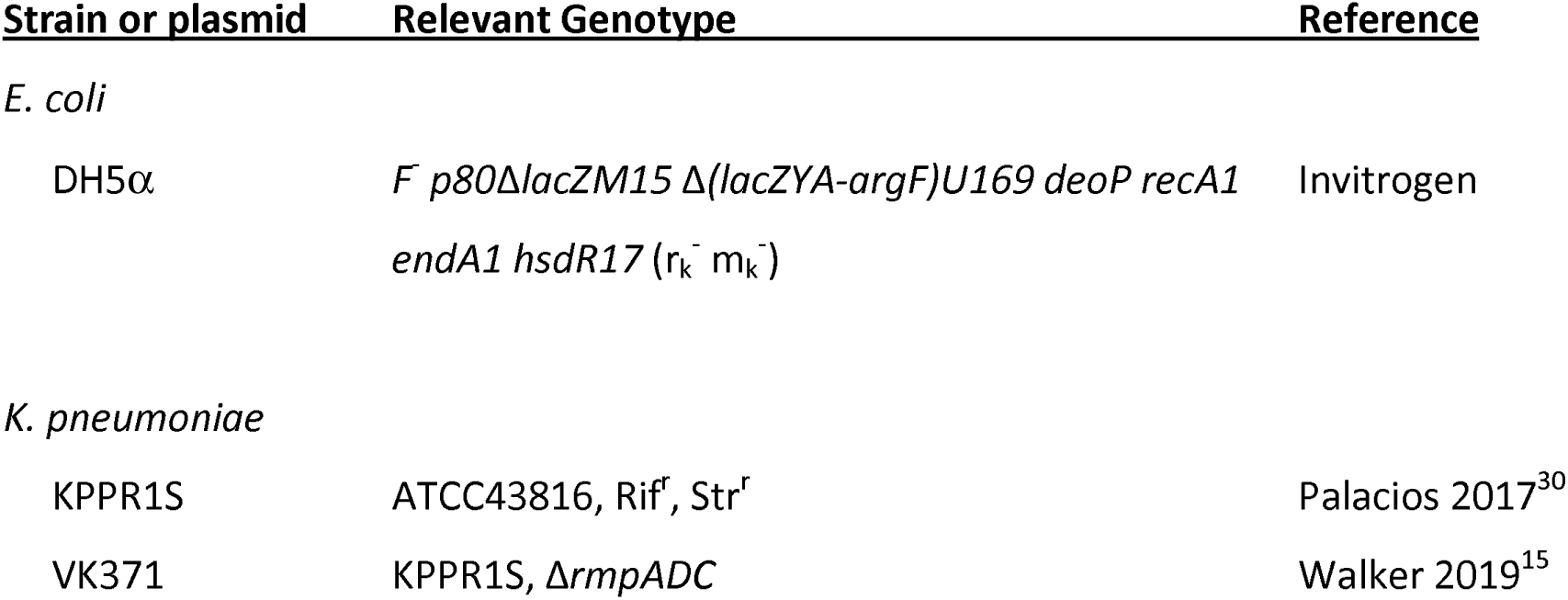
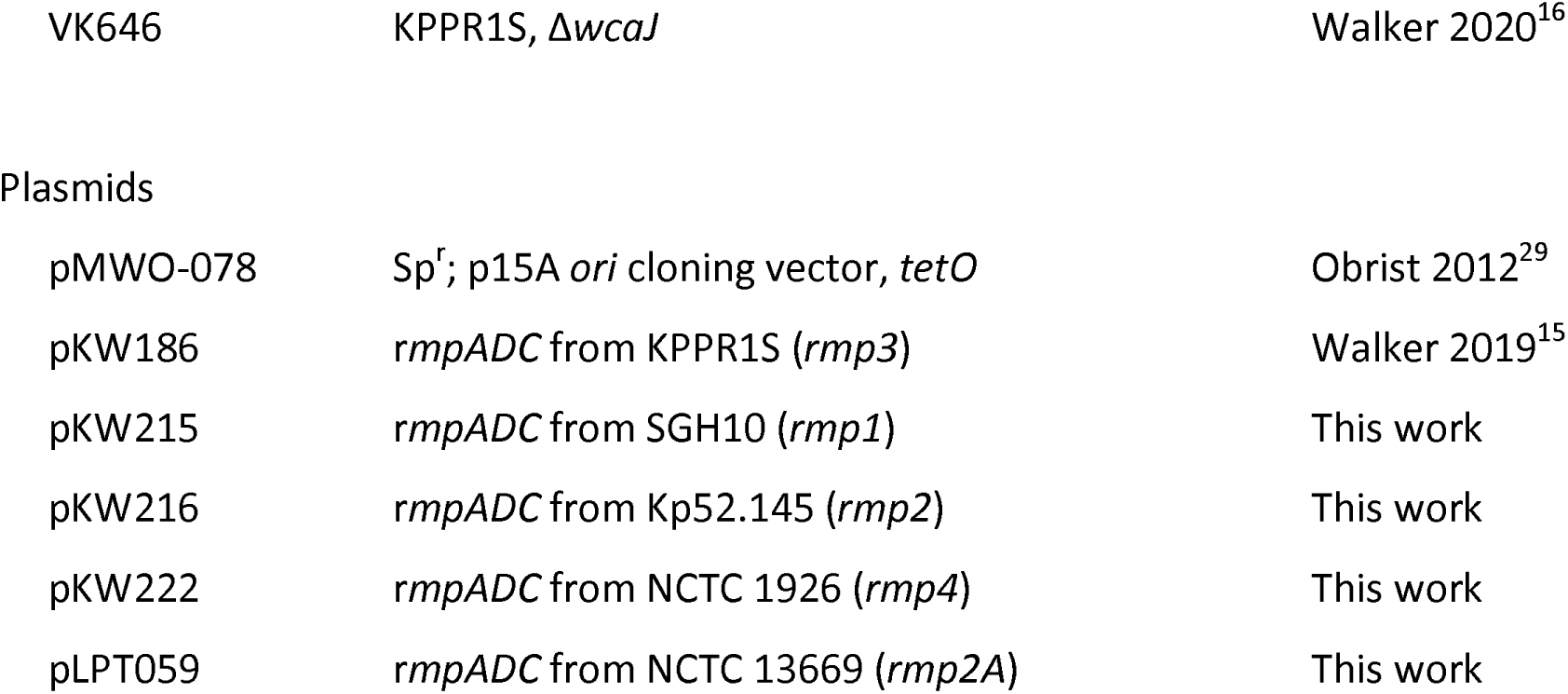
Strains and plasmids used in this work.

Primers used for the construction of expression vectors are listed in **Additional File 2**. All *rmp* loci were amplified from genomic DNA by PCR and then cloned into pMWO-078^29^ by Gibson assembly (NEB). The *rmp* expression vectors were then transformed into KPPR1S Δ*rmp*. KPPR1S (ST493; K2/O1) is a well-characterised streptomycin and rifampicin-resistant derivative of ATCC 43816, and has been used in previous studies to characterise *rmpD, rmpC*, and HMV^15–17^. Wildtype KPPR1S, Δ*wcaJ*, and Δ*rmp* strains transformed with pMWO-078 were also included as control strains for measuring HMV. The *wcaJ* mutant does not produce capsule and is HMV negative, while the *rmp* mutant produces capsule but is HMV negative as it does not produce RmpD.

### Assessment of hypermucoviscosity

Saturated overnight cultures of strains containing *rmp* plasmids were subcultured in fresh LB for 5.5 hours at 37°C with 100 ng/ml anhydrotetracycline (ATc) to induce *rmp* expression. Cultures were normalised to 1 OD_600_/ml and centrifuged at 1000 x *g* for 5 minutes. Mucoviscosity of cultures was determined by normalising OD_600_ of the culture supernatant to the starting culture as previously described^31^.

### Uronic acid measurement

Uronic acid (UA) was measured following an established protocol^32^ from cultures grown as described for assessing HMV. Briefly, UA was extracted from 500 μl of culture with zwittergent, precipitated with ethanol, and resuspended in tetraborate/sulfuric acid. Phenylphenol was added and absorbance at 520 nm was measured. UA amounts were determined from a standard curve generated with glucuronolactone.

## Results

### Prevalence of *rmp* in the *K. pneumoniae* species complex

Screening for the *rmp* locus in publicly available KpSC genomes identified its presence in 992/13993 genomes across three KpSC species; *K. pneumoniae* (980/11967, 8.2%), *K. quasipneumoniae* subsp. *similipneumoniae* (*Kqs*; 5/522, 1.0%) and *K. variicola* subsp. *variicola* (*Kv*; 7/626, 1.1%) (see **Table 2**). The majority of these included complete *rmp* loci with intact coding sequences for *rmpA*, *rmpC* and *rmpD* (i.e. functional variants, detected in 73% of *rmp*+ genomes), however some loci were incomplete (i.e. deletion variants missing at least one gene within the locus or arising from assembly fragmentation) or had mutations resulting in premature stop codons truncating one or more of the encoded proteins (i.e. truncation variants) (see **Table 2**). Fifteen genomes were identified with multiple *rmp* loci (two *rmp* each); the majority of these carried at least one functional *rmp* locus (73%; eight genomes with two functional *rmp* loci and three genomes with one functional *rmp*) while the remaining four genomes each had two non-functional *rmp* loci arising from deletion and/or truncation variants. BLASTn search of all non-redundant bacteria in NCBI with default parameters identified complete *rmp* loci in four non-*Klebsiella* genomes, in all cases *rmp* was present in plasmids sequenced from *Escherichia coli* transconjugants that had been mated with a *K. pneumoniae* strain carrying KpVP-1-like virulence plasmids (GenBank accessions MN200130.1, MN182750.1, MZ475697.1, CP068571.1). Incomplete *rmp* loci (44% coverage) were detected in an additional five plasmids that were also sequenced from *E. coli* transconjugants.

**Table 2.** Distribution of *rmp* loci by species.

|  | *K. pneumoniae* | *K. quasipneumoniae*<br>subsp.<br>*similipneumoniae* | *K. variicola*<br>subsp.<br>*variicola* |
| --- | --- | --- | --- |
| Number of genomes screened | 11967 | 522 | 626 |
| ***Rmp* status** |  |  |  |
| Any *rmp* sequence detected | 980 | 5 | 7 |
| i. Functional *rmp* | 709 | 3 | 4 |
| ii. Incomplete *rmp* | 115 | 1 | 3 |
| iii. Truncated *rmp* | 141 | 1 | - |
| iv. Multiple *rmp* | 15 | - | - |

### Genetic diversity of the *rmp* locus

Each of the genes comprising the *rmp* locus showed some degree of genetic diversity, with 85 *rmpA* alleles, 77 *rmpD* alleles and 42 *rmpC* alleles observed across *rmp*+ genomes. For the 825 genomes carrying an intact ‘typeable’ *rmp* locus (838 loci total accounting for genomes with two *rmp* loci), allelic variants could be assigned to all three genes, and resulted in 170 unique combinations which were each assigned a unique RmST (*rmp* sequence type). The *rmp* loci from the remaining 167 genomes were designated ‘non-typeable’ due to the locus being incomplete (i.e. missing at least one gene or comprising a fragmented gene to which an allele could not be designated).

Maximum likelihood phylogenetic analysis of the concatenated *rmpA*, *rmpD* and *rmpC* sequences belonging to each unique RmST revealed the grouping of sequences into five distinct lineages, which we labelled *rmp1, rmp2, rmp2A, rmp3* and *rmp4* (**Figure 1**). These lineages were labelled as such to match with their associated *iro* and/or *iuc* lineages. The nucleotide divergence between lineages ranged from 0.7-11% (mean 4.7%), decreasing to 0-5.9% within lineages (mean 0.4%). No *rmp* gene alleles were shared between lineages (see individual gene phylogenies in **Additional File 3**), suggesting an absence of recombination between the lineages. The *rmp3* lineage was notably divergent from the other *rmp* lineages, with a mean divergence of 8.4% compared to 1.8% between all other lineages (**Table 3**). Further, the *rmpD* alleles of the *rmp3* lineage were longer than those of other lineages, measuring 176-177 bp in length compared to the 151-162 bp alleles observed in *rmpD* from the other lineages.

**Figure 1.**
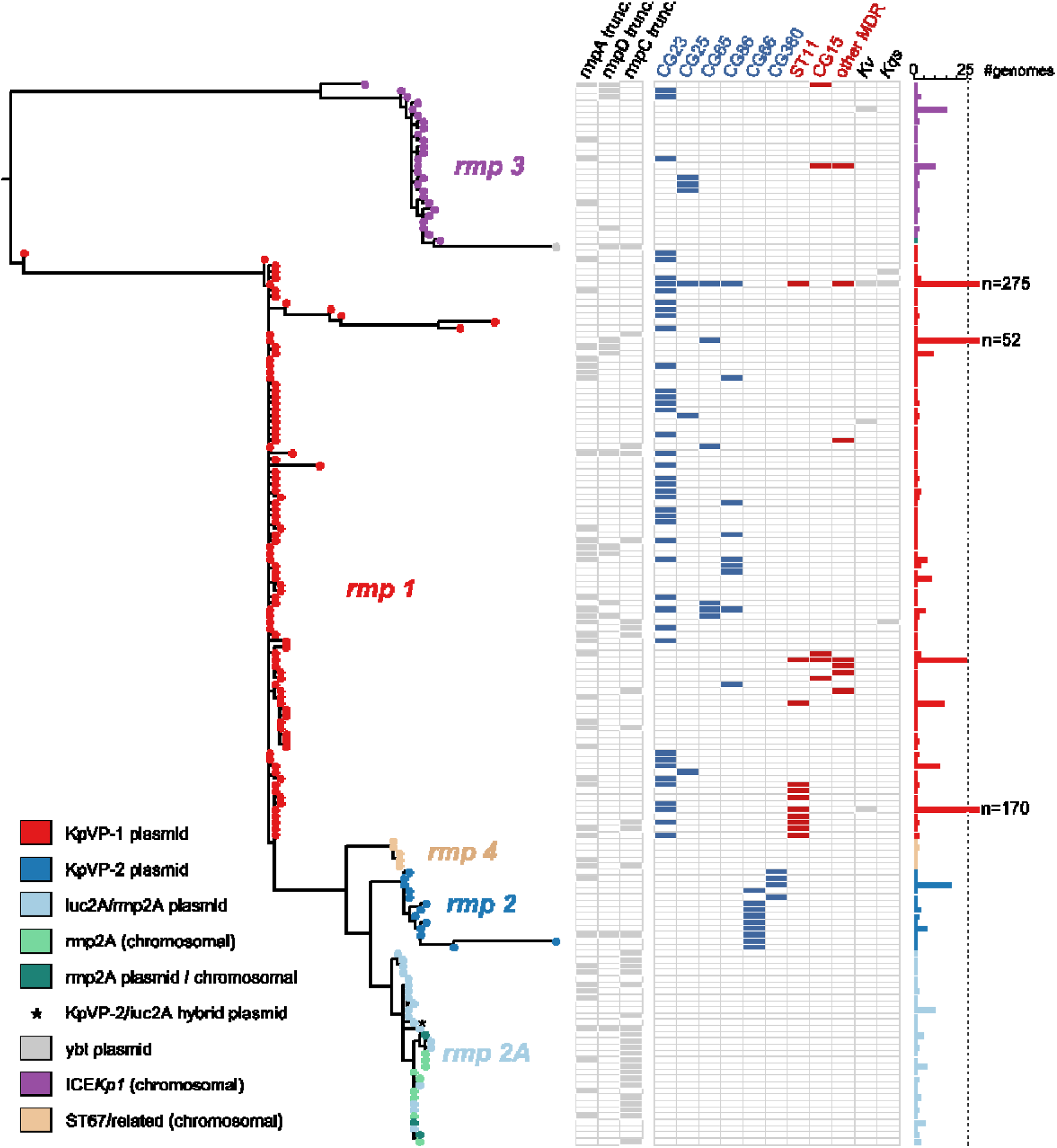
Maximum-likelihood phylogenetic tree of *rmp* sequence types (RmSTs). Lineages are labelled and tips are coloured by the associated mobile genetic element according to the legend. Columns are as follows for a particular RmST: presence or absence of truncations in the *rmpA*, *rmpD* and *rmpC*, detection within a hypervirulent (blue) or MDR (red) clone or non-*K. pneumoniae* species (Kv, *Klebsiella variicola*; Kqs, *Klebsiella quasipneumoniae* subsp. *similipneumoniae*). The number of genomes from which each RmST was detected is shown in the bar graph on the right-hand side.

**Table 3.**
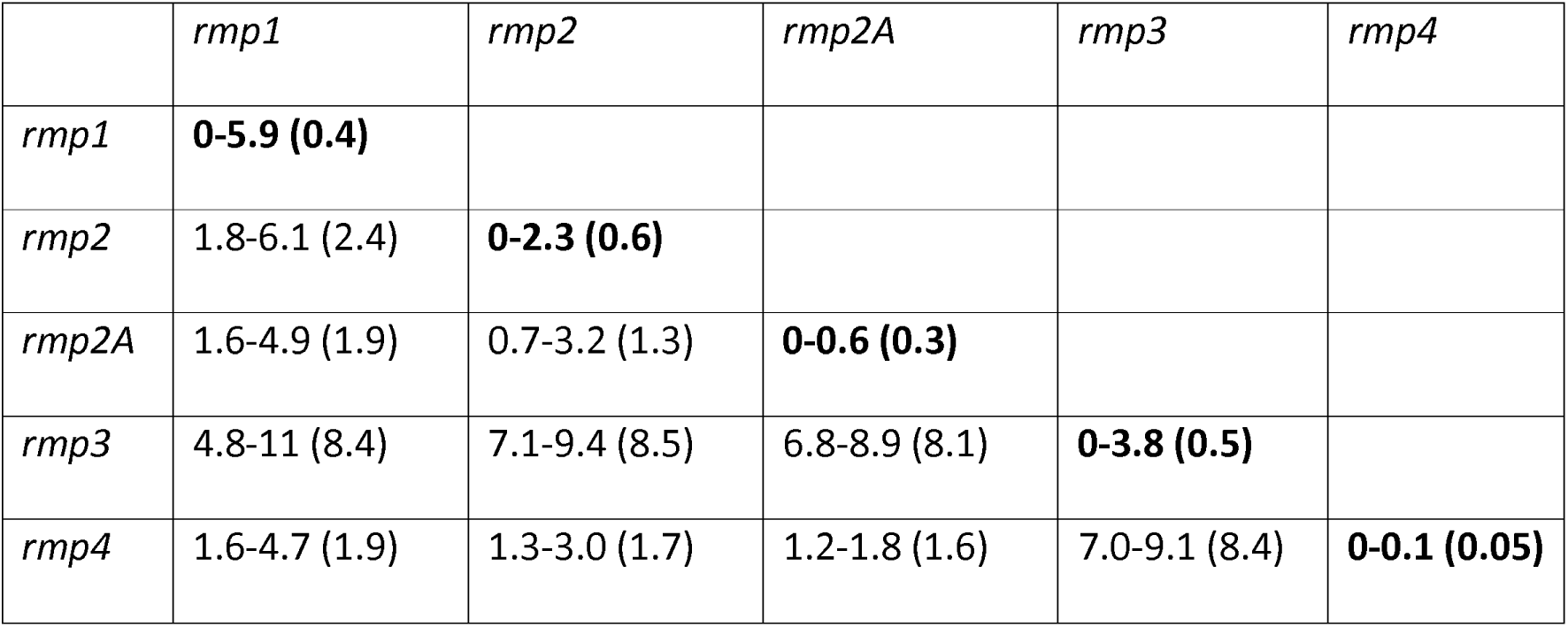
Nucleotide divergence within and between *rmp* lineages. Values given are range (mean) and expressed as percentages.

|  | *rmp1* | *rmp2* | *rmp2A* | *rmp3* | *rmp4* |
| --- | --- | --- | --- | --- | --- |
| *rmp1* | **0-5.9 (0.4)** |  |  |  |  |
| *rmp2* | 1.8-6.1 (2.4) | **0-2.3 (0.6)** |  |  |  |
| *rmp2A* | 1.6-4.9 (1.9) | 0.7-3.2 (1.3) | **0-0.6 (0.3)** |  |  |
| *rmp3* | 4.8-11 (8.4) | 7.1-9.4 (8.5) | 6.8-8.9 (8.1) | **0-3.8 (0.5)** |  |
| *rmp4* | 1.6-4.7 (1.9) | 1.3-3.0 (1.7) | 1.2-1.8 (1.6) | 7.0-9.1 (8.4) | **0-0.1 (0.05)** |

### Diversity of *rmp-*associated mobile genetic elements

Each of the *rmp* lineages were associated with unique plasmids or chromosomal contexts (see **Figure 1**), many of which have been previously characterised in a study examining the aerobactin- and salmochelin- encoding loci *iuc* and *iro*^10^, which are typically co-localised with *rmpA2* and *rmp* respectively on the same mobile elements. The dominant *K. pneumoniae* virulence plasmid, KpVP-1 (associated with *iuc1* and *iro1* loci) was associated with the *rmp1* lineage and accounted for the majority of *rmp* carriage (785 genomes, 79.1%). Next most common were the *iuc2A* virulence plasmids (associated with *iuc2A*)^10^, carrying *rmp2A* lineage (77 genomes, 7.8%); followed by ICE*Kp1* carrying *rmp3* (58 genomes, 5.8%); and KpVP-2 virulence plasmids with *rmp2* (associated with *iuc2* and *iro2*, 45 genomes, 4.6%). Two additional genetic contexts were observed in this study: (i) the *rmp4* lineage was chromosomally-encoded and associated exclusively with *K. pneumoniae* CG67 genomes (six genomes, identified as ST67 or single-locus variants thereof, also known as *K. pneumoniae* subspecies rhinoscleromatis^33^); and (ii) transposition of *rmp3* into the yersiniabactin-encoding *ybt4-*type plasmids (three genomes; note partial deletion of *rmp* locus in two of these genomes). For the remaining genomes, one was associated with a KpVP-2/*iuc2A* hybrid plasmid as previously described^10^, nine carried multiple *rmp* loci (due to presence of both KpVP-1 and ICE*Kp1*), and 17 were unresolved due to assembly issues. KpVP-1 was detected in 785 genomes, where it typically carried both *rmp* and *rmpA2* (669/785). However, deletion variants of KpVP-1 were common (i.e. coverage of KpVP-1 reference ranged from 26.7% coverage in *rmp1*+ genomes): we observed 116/785 genomes with *rmp* only without *rmpA2* (20 and 32 of these lacking an intact *iro1* and *iuc1*, respectively).

The genetic context of *rmp* in representatives of each of the key *rmp* lineages is shown in **Figure 2**. The *rmp* locus was located adjacent to the *iro* locus and *peg-344* gene (which has been mis-labelled as *pagO* in some studies) in most cases, with the exception of the *ybt4* plasmids (which lack *<u>iro</u>* and *peg-344* genes) and *iuc2A* plasmids (which carry *iroB* only). The *iro* locus is typically intact in KpVP-1, KpVP-2 and ICE*Kp1,* but is partially disrupted by insertion sequences (IS) in the ST67 (rhinoscleromatis) chromosome and *iuc2A* plasmids. IS flank the *rmp*/*iro* region in all contexts, however the specific IS vary and it is not clear which have played a role in mobilising *rmp* and/or *iro*. Notably, all contexts include IS*3* on one or both ends of the *rmp*/*iro* region. The closely related *rmp1*, *rmp2, rmp2A* and *rmp4* lineages all have IS*3* in a conserved position adjacent to *rmpA*; this also appears to be conserved in the divergent lineage *rmp3,* suggesting it was likely present in the common ancestor of all *rmp* lineages and may have played a role historically in the mobilisation of *rmp*/*iro* between genetic backgrounds (see **Figure 2**). However all *rmp/iro* regions have additional IS near or within the locus, some of which may have contributed to further mobilisation and/or degradation of the distinct lineages over time. While the assembly of the *rmp* region in the *ybt4* plasmid was incomplete, an IS*3* fragment was detected next to *rmp*.

**Figure 2.**
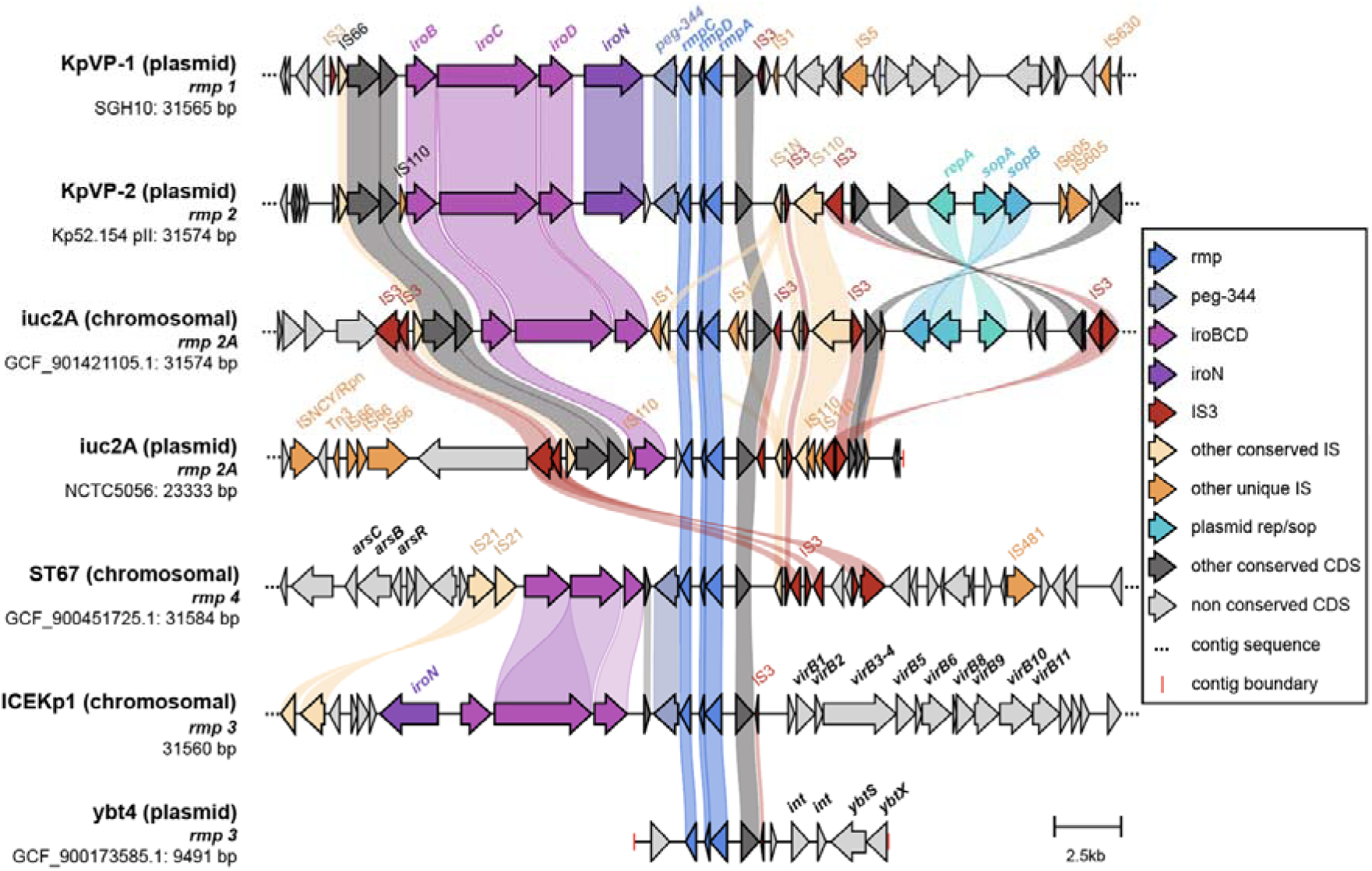
Comparison of upstream and downstream regions of *rmp* associated with different mobile variants. Arrows represent coding sequences and those corresponding to genes of interest are labelled and coloured by functionality as per the legend. Other labelled loci include *arsCBR* (arsenic resistance), *virB1*-*virB11* (*virB*-type 4 secretion system) and *ybtSX* (part of the yersiniabactin locus). Shading correspond to regions of similarity (sequence identity ≥ 30%) as identified by clinker.

### Hypermucoviscosity of *rmp* variants in a KPPR1S background

To determine if there were functional differences between the level of HMV conferred by the *rmp* lineages, we ectopically expressed variants of the *rmp* locus cloned into pMWO-078 in an HMV-negative *rmp* deletion mutant KPPR1S Δ*rmp*. Wildtype KPPR1S is a ST493 K2/O1 isolate containing *rmp3* (ICE*Kp1*), which is the most divergent *rmp* lineage (**Figure 1**); KPPR1S Δ*rmp* therefore provides a good background to assess potential functional differences between *rmp* variants. HMV was assessed using a sedimentation assay. The Δ*wcaJ* and Δ*rmp* vector controls fully sedimented, whereas the KPPR1S vector control did not sediment well (**Figure 3A**). Ectopical expression of the native locus, *rmp3,* conferred elevated HMV in Δ*rmp* compared to the KPPR1S vector control. This elevated phenotype is likely due to the overexpression of *rmp,* and this complemented strain therefore serves as a reference to compare the effects of expressing the other *rmp* loci. Expression of the *rmp* genes from *rmp1, rmp2 and rmp2A* also conferred elevated HMV levels in Δ*rmp* (**Figure 3A**), and there were no statistically significant differences in sedimentation resistance between these strains versus that expressing *rmp3*. However, expression of *rmp4* conferred a significantly lower level of HMV compared to the other lineages.

**Figure 3.**
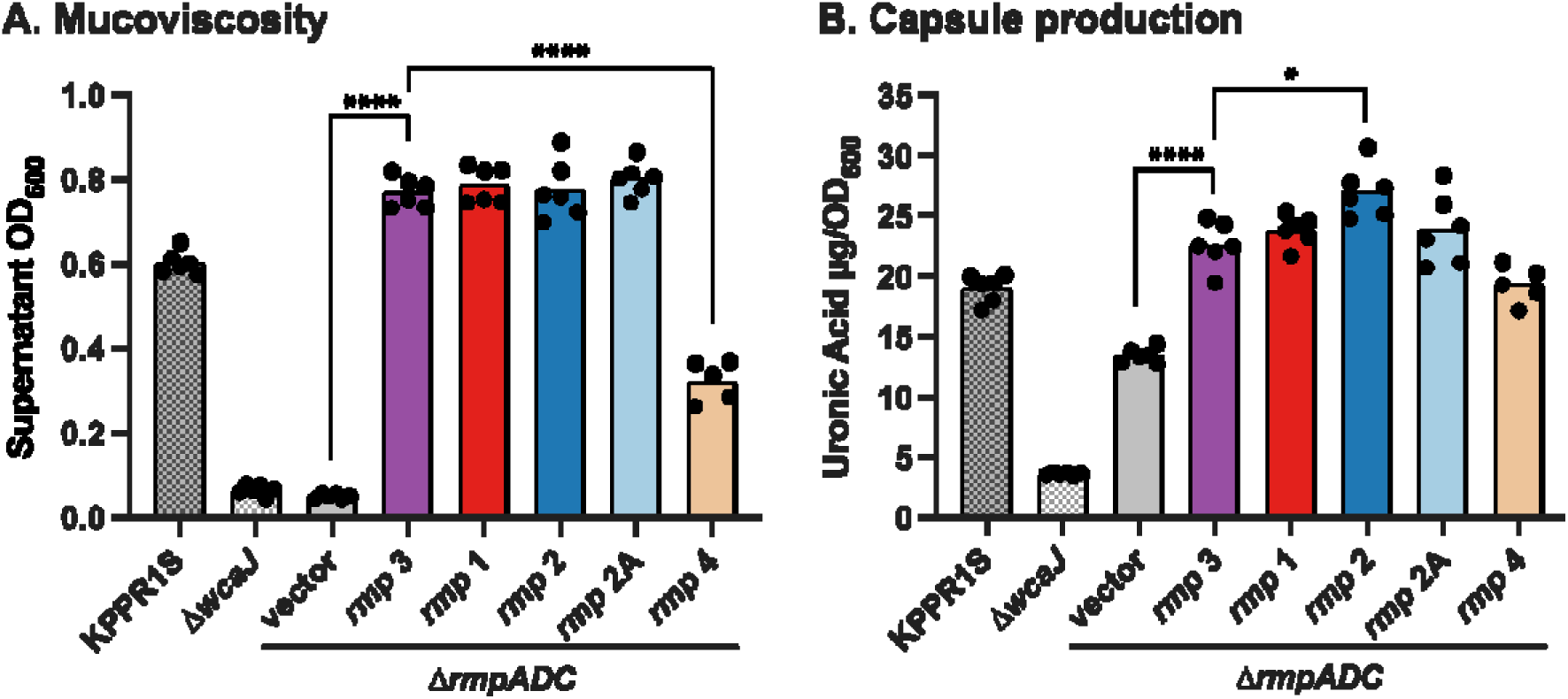
Hypermucoviscosity and capsule production of ectopically expressed *rmp* loci from the five different lineages. A. Mucoviscosity assay, and B. uronic acid assay of KPPR1S, Δ*wcaJ*, and Δ*rmp* strains with vector (pMWO-078) or lineage-specific pRmp (see Table 1). Data from two technical replicates and three biological replicates were obtained following a 5.5 hour induction of KPPR1S expression plasmid-borne *rmp* genes as described in Methods. One-way ANOVA with Tukey’s post-test was used to determine significance. *, *P* < 0.05; ***, *P* < 0.001; ****, *P* < 0.0001; comparisons not indicated were not significant.

### Capsule production of *rmp* variants in a KPPR1S background

Elevated capsule production is an *rmpC*-dependent phenotype in KPPR1S, independent of HMV, whereby RmpC upregulates capsule expression via the promoters upstream of the *galF* and *manC* genes in the K locus, and KPPR1S Δ*rmp* mutants display decreased capsule production^15,16^. To further investigate any functional differences between *rmp* variants, we assayed the same transformants described above for uronic acid (UA) production, which is used as an indicator of the amount of capsule^32^. KPPR1S, Δ*wcaJ* (i.e. capsule-negative), and *Δrmp* strains were included as controls. Expression of *rmp* from each of the lineages increased capsule production (**Figure 3B**). Expression of *rmp1, rmp2A* and *rmp4* in the *Δrmp* mutant increased capsule production to the same level observed when expressing the native *rmp3* locus (**Figure 3B**). Expression of *rmp2* produced slightly but significantly higher uronic acid levels compared to *rmp3* (p < 0.05). Collectively, these data suggest that there are subtle but significant functional differences between the *rmp* lineages.

### Distribution of *rmp* in the *K. pneumoniae* species complex

The *rmp* locus was detected in 129 unique *K. pneumoniae* STs, 3 *Kqs* STs and 5 *Kv* STs, representing 143 unique combinations of STs and *rmp* lineages. As expected, the prevalence of *rmp* is typically quite high within known hvKp clones (≥80% prevalence within any given ST assigned to a hypervirulent clonal group with the exception of CG25, which has recently been shown to comprise of two distinct lineages; mean 83.6%) (**Figure 4**). *Rmp* was also common in CG67 (rhinoscleromatis, 100% *rmp4*; 25% truncated) and CG91 (ozaenae, 79% *rmp2A* incomplete or truncated); both lineages have previously been defined as subspecies due to their distinct pathotypes^33^. However, relatively high *rmp* prevalence was also observed in some ‘generalist’ (i.e. non-HvKp and non-MDR) *K. pneumoniae* clones (1.6-100% *rmp* prevalence, mean *rmp* prevalence=75.0%). Conversely, the MDR clones together with CG36 and CG45, which are also clones that are commonly detected in healthcare settings, had relatively lower frequencies of *rmp* (mean prevalence=11.4%).

**Figure 4.**
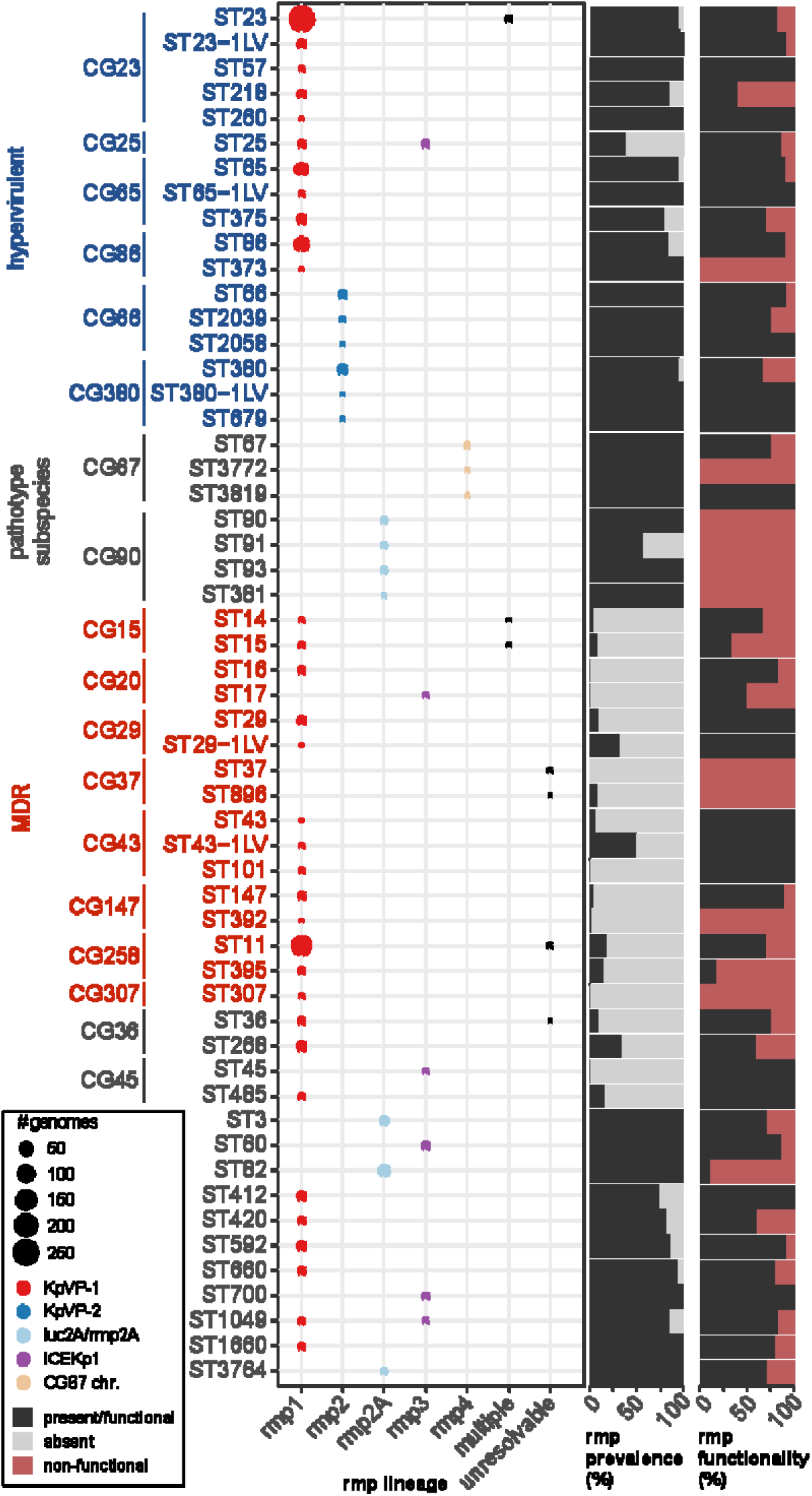
Distribution of *rmp* and *rmpA2* across *K. pneumoniae* sequence types with ≥5 *rmp+* genomes. Rows indicated *K. pneumoniae* sequence types (STs; as labelled), and grouped and labelled by clonal group (CG) where applicable; those corresponding to hypervirulent, pathotype subspecies or multidrug-resistant (MDR) clones are labelled accordingly. The bubble plot shows the number of *rmp*+ genomes assigned to a particular *rmp* lineage, coloured by the type of mobile element. The first bar plot shows prevalence of *rmp* (i.e. black: present, grey: absent) and the second shows functional status of the locus (i.e. black: intact and functional, red: non-functional due to deletions or truncations).

KpVP-1-*rmp1* was the most widely disseminated, and was not only detected in the hypervirulent clones from which they were initially characterised but also in MDR clones. Numbers of KpVP-1-*rmp1* were notably high in ST11 (211 genomes) and ST15 (50 genomes), reflecting recently reported MDR-hypervirulent convergent variants of these well-known MDR clones^34^. KpVP-1 also accounted for the majority of *rmp* acquisitions in the other KpSC species where the genetic context could be resolved. In comparison, the other mobile elements appeared in relatively fewer STs, but also appeared to be fixed in most of these clones that likely serve as native hosts to the respective mobile elements (see **Figure 4**).

Amongst *rmp*+ genomes with a confident K locus call (917/992), 38 different K loci were detected (**Figure 5**). Half (i.e. 19) were detected in two or fewer genomes. The most common K loci were KL1 (319 genomes), KL2 (203), KL64 (140), which collectively account for 72.2% of genomes with a confident K locus, followed by KL57 (45) and KL20 (40). The dominant K loci are largely driven by their associations with over-represented STs (**Figure 5**), many of which are known hvKp lineages such as KL1 in CG23 (which harbours KpVP-1), and KL2 in CG380 (KpVP2), CG65 and CG86 (KpVP-1). The next most common K locus, KL64, is associated with a Chinese sublineage of MDR clone ST11, known to have acquired variants of KpVP-1^34^.

**Figure 5.**
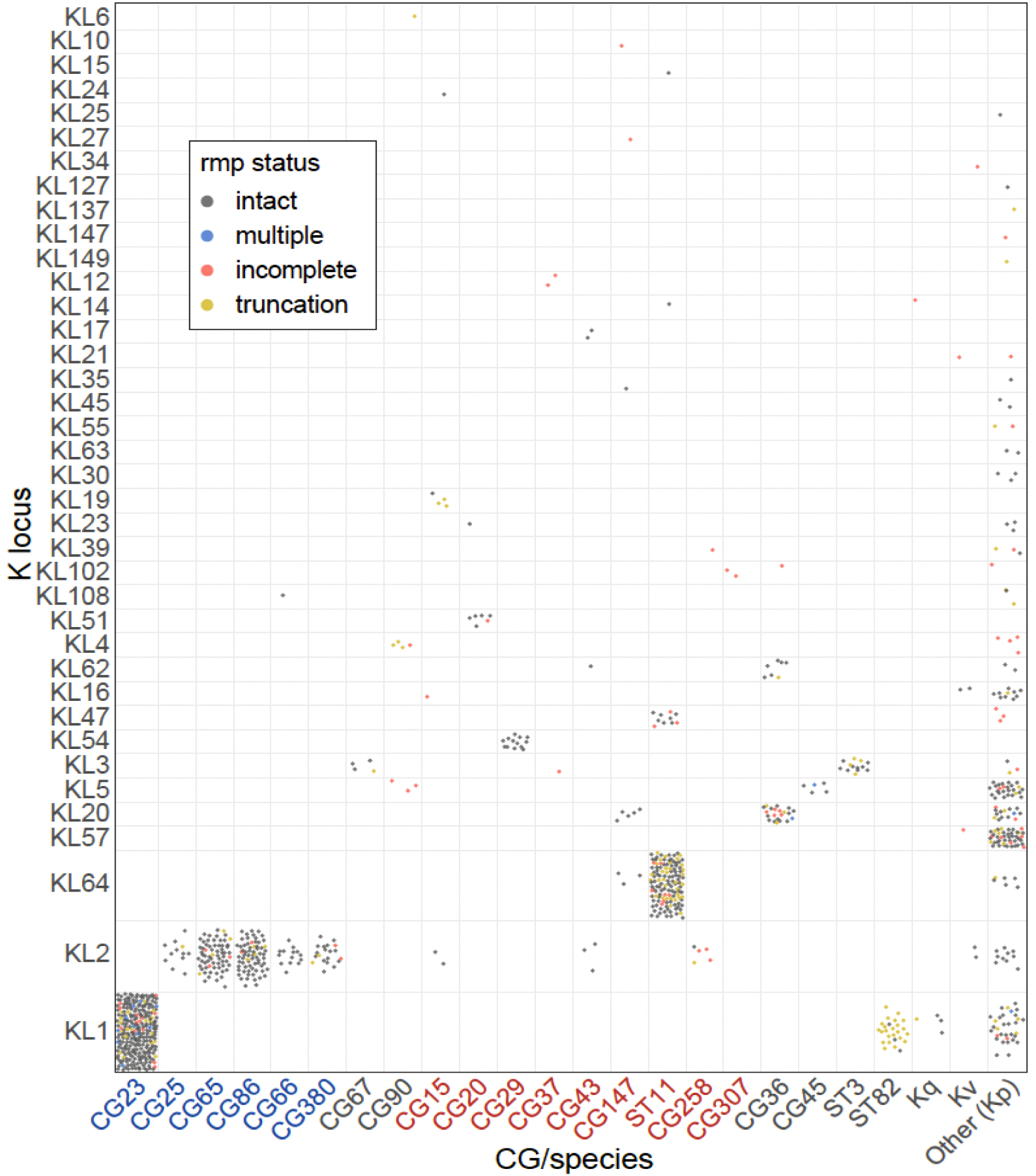
Distribution of *rmp* within K locus groups and KpSC clonal groups (CG) or species. Each circle represents an *rmp*+ genome with a confident K locus call (917 genomes), and is coloured by the functionality of the *rmp* locus as per the figure legend. Clonal group labels are coloured blue and red to indicate hypervirulent and MDR lineages, respectively. Kq: *Klebsiella quasipneumoniae*, Kv: *Klebsiella variicola*.

Given previous reports and observations highlighting the common occurrence of truncations in *rmpA* due to insertions and deletions (indels) within a poly-G tract^14^, we next examined the *rmp* locus for the occurrence of truncations. Truncations were detected in 30/85 *rmpA*, 9/77 *rmpD* and 16/42 *rmpC* alleles across 63, 16 and 55 genomes respectively (**Additional File 3**). The majority of truncations (45/55 truncated alleles) were caused by indels within homopolymer tracts resulting in premature stop codons arising from frameshift mutations (**Additional File 4**). Further, some homopolymer regions had a higher frequency of mutations compared to others, including the previously reported poly-G tract in *rmpA* spanning nucleotide position 276 to 285 whereby indels were detected in 16 *rmpA* variants across 46 genomes. The frequency of truncated *rmp* loci compared to intact loci was lower within the hypervirulent clones and appeared to be more common outside of these clones (**Figures 4** and **5**). Closer inspection of the phylogenies for hypervirulent clones CG23, CG65 and CG86 (**Additional Files 5 - 7**) revealed a random distribution of truncated *rmp* alleles throughout these populations.

## Discussion

We previously reported briefly on the sequence variation of the *rmp* locus, which clustered into four lineages and were each associated with distinct MGEs^20^. In this study we provide detailed insights into the phylogenetic relationships, genetic contexts, distribution and phenotypic functionality of *rmp* variants. Building on the initial *rmp* scheme, we characterised an additional *rmp4* lineage observed only in genomes belonging to a sublineage of *K. pneumoniae* (CG67), updating the number of *rmp* lineages to five.

The 1:1 correlation observed between *rmp* lineages and the MGEs that mobilise them also extends to *iro,* shown here to be typically located adjacent to *rmp* (**Figure 2**), and to *iuc,* which is often located elsewhere on the same plasmids^10^. This association highlights the co-evolution of these virulence loci, and presumably other common genes shared between the MGEs. While specific details or the order of events in the evolutionary history of these MGEs cannot be determined, similarities in the genetic contexts surrounding *rmp* do suggest a shared ancestry and the role of IS, particularly IS*3*, in the mobilisation of *rmp/iro* between different MGEs. Interestingly, the presence of IS*3* upstream of *rmpA2* has been flagged as being necessary for complete activation of the promoter for *rmpA2*, and may likewise serve a similar purpose for *rmpA*^35^. We also observed three instances of a novel genetic context for *rmp*, where a variant of *rmp3* (ICE*Kp1*) had been introduced into a yersiniabactin plasmid (i.e. another virulence MGE)^36^ likely via IS*3*-mediated transposition (**Figure 2**). Outside of *K. pneumoniae*, *rmp* (KpVP-1 and ICE*Kp1*) was only detected in two other KpSC species at rare frequencies, and did not appear to have been acquired naturally in any other species. *K. pneumoniae* is therefore likely to be the original host for *rmp,* although the same may not hold true for the other virulence loci given that *iro* and *iuc* variants have been detected in other non-*Klebsiella* species including *Enterobacter spp.* and *Escherichia coli*^10^.

Even with the larger genome dataset screened in this present study, the prevalence of *rmp* overall (7.5%) and each of the MGEs was very similar to that reported in our earlier study characterising *iuc* (8.7%) and *iro* (7.2%) (i.e. 13000 versus 2700 genomes). The *rmp1* lineage (KpVP-1) was by far the most dominant accounting for 80% of the *rmp* burden, in part driven by their association and maintenance via clonal expansion within the key hypervirulent clones such as CG23^11^, but also due to the more recent transmission and spread within MDR clones such as ST11 and ST15 (**Figures 1** and **4**). Based on growing reports of ‘hypervirulent ST11’ from China, it is possible that KpVP-1 is also being stably maintained within this clone following its acquisition. With the exception of KpVP-1-*rmp1* and ICEKp1-*rmp3*, which were detected in multiple STs/CGs, the remaining MGEs also appeared to be stably conserved within a small number of clones in which they were detected (93.9-100% prevalence; **Figure 4**).

Each of the *rmp* lineages was shown to be functional, resulting in HMV and elevated capsule production when a representative of each was introduced into a single strain background, KPPR1S *rmp*. Notably, this ST493 strain has a K2 capsule, which is one of two dominant capsule types among known hvKp clones such as CG86 (**Figure 5**). We also observed variability in the extent of HMV and capsule production for different *rmp,* which further reiterates that HMV and capsule production are two separable traits^15,16^; for example, the expression of *rmp4* does not restore HMV to the same level as KPPR1S *rmp3* but does restore capsule production (**Figure 3**). It is unclear if expressing these *rmp* (or other representatives of the same lineages) in other KpSC strains with different serotypes will have the same impact, especially given the extensive diversity of Wzc in different K loci and its interaction with RmpD. These insights will be particularly useful for predicting the impacts of *rmp* acquisition in other clones, including those considered to be MDR, and is the subject of ongoing work by our team.

Loss of function mutations of *rmpA* arising from indels within the poly(G) tract has been well documented in many studies^14^, and while this site (i.e. bases 267 to 285 of the *rmpA_2* reference) does account for the majority of *rmpA* truncations in this dataset, indels within additional homopolymer tracts in *rmpA, rmpD* and *rmpC* were also observed (**Additional File 4**). Loss of function mutations in the *rmp* genes were observed in at least 145 genomes (14.6% of *rmp+* genomes, including three with multiple *rmp*), most often in the non-hypervirulent clones. Further, we observed parallel evolution of the same loss of function mutations on multiple occasions within different hypervirulent clones (e.g. allele *rmpA_4* in CG23, CG65 and CG86; **Additional Files 5-7**). Taken together, these findings highlight the potential reversibility of homopolymer tract mutations, which may take place to help alleviate the negative selection pressure following acquisition of KpVP-1 or other MGEs. Importantly, these loss of function mutations also need to be carefully considered when interpreting data based solely on PCR detection of *rmpA/A2*, and may partly explain the discrepancies in the literature reporting on the association between *rmp* presence and HMV. Although, other reasons also include inconsistencies and the unreliability of string testing, which is heavily influenced by temperature dependencies and strain genetic background^37^.

## Conclusions

Our findings reveal that, similar to the other key virulence loci co-localised on the same MGEs, genetic variation within the *rmp* locus is highly structured and this information can be harnessed to track novel *rmp* acquisitions. To this end, detection and genotyping of the *rmp* genes (i.e. RmST typing), alongside the reporting of locus disruptions, has already been implemented in our genotyping tool for KpSC genomes, Kleborate (github.com/klebgenomics/Kleborate). The tool also outputs a virulence score that is currently calculated from the presence of various siderophores (*ybt* and iuc) and colibactin (*clb*), and does not take into account *rmp* (or *iro*). Given the apparent role of *rmp* in hypervirulence, the virulence score in addition to *rmp* presence should both be considered when assessing the virulence of a given strain or genome. Ongoing research investigating the expression of *rmp* variants in different strain and capsule backgrounds will yield additional important insights into the function of *rmp,* particularly for novel strains such as those from the MDR clones that acquire them, and can be used to further improve the genotyping output for *rmp*.

## Declarations

### Ethics approval and consent to participate

Not applicable

### Consent for publication

Not applicable

### Availability of data and materials

All whole-genome sequences analysed in this study are publicly available on NCBI, and the accession numbers are listed in Additional File 1. The RmST scheme is available in the *K. pneumoniae* BIGSdb database (https://bigsdb.pasteur.fr/klebsiella/) and in the Kleborate distribution (https://github.com/klebgenomics/Kleborate)

### Authors’ contributions

Acquisition of data: MMCL, SMS, LPT, LMJ, KAW

Analysis and interpretation of data: MMCL, SMS, LPT, KLW, SB, KAW, VLM, KEH

Writing of code: RRW and KEH

Initial drafting of manuscript: MMCL with input from KEH and KLW

Conceptualisation of study: KEH with input from VLM and KAW

All authors edited and approved the submitted manuscript.

## Acknowledgements

We thank the Institut Pasteur curation team of BIGSdb-Pasteur for importing novel alleles, profiles and/or isolates at https://bigsdb.pasteur.fr/.

---

**Additional file 1**

Strain information and Kleborate genotyping output for genomes included in this study (XLS)

**Additional file 2**

Primers used in this study (DOC)

**Additional file 3**

Phylogenetic relationships of the A. *rmpA,* B. *rmpD* and C. *rmpC* genes. (DOC)

**Additional file 4**

Description of truncated allelic variants of *rmpA, rmpD* and *rmpC* (XLS)

**Additional file 5**

Distribution of *rmpADC* allelic variants and KpVP-1 associated virulence loci *rmpA2*, *iuc* and *iro* in *Klebsiella pneumoniae* clonal group 23 (DOC)

**Additional file 6**

Distribution of *rmpADC* allelic variants and KpVP-1 associated virulence loci *rmpA2*, *iuc* and *iro* in *Klebsiella pneumoniae* clonal group 65 (DOC)

**Additional file 7**

Distribution of *rmpADC* allelic variants and KpVP-1 associated virulence loci *rmpA2*, *iuc* and *iro* in *Klebsiella pneumoniae* clonal group 86 (DOC)

## References

1. Podschun, R. & Ullmann, U. *Klebsiella* spp. as Nosocomial Pathogens: Epidemiology, Taxonomy, Typing Methods, and Pathogenicity Factors. Clin Microbiol Rev 11, 589–603 (1998).

2. Shon, A. S., Bajwa, R. P. S. & Russo, T. A. Hypervirulent (hypermucoviscous) *Klebsiella pneumoniae*: a new and dangerous breed. Virulence 4, 107–118 (2013).

3. Russo, T. A. & Marr, C. M. Hypervirulent *Klebsiella pneumoniae*. Clin. Microbiol. Rev. 32, e00001–19 (2019).

4. Lee, I. R. et al. Differential host susceptibility and bacterial virulence factors driving *Klebsiella* liver abscess in an ethnically diverse population. Sci Rep 6, 29316 (2016).

5. Choby, J. E., Howard-Anderson, J. & Weiss, D. S. Hypervirulent *Klebsiella pneumoniae* – clinical and molecular perspectives. J. Intern. Med. 287, 283–300 (2020).

6. Struve, C. et al. Mapping the evolution of hypervirulent *Klebsiella pneumoniae*. MBio 6, 1–12 (2015).

7. Wyres, K. L., Lam, M. M. C. & Holt, K. E. Population genomics of *Klebsiella pneumoniae*. Nat. Rev. Microbiol. 18, 344–359 (2020).

8. Lam, M. M. C., Wick, R. R., Judd, L. M., Holt, K. E. & Wyres, K. L. Kaptive 2.0: updated capsule and lipopolysaccharide locus typing for the *Klebsiella pneumoniae* species complex. *Microb*. Genomics 8, (2022).

9. Lin, T. L., Lee, C. Z., Hsieh, P. F., Tsai, S. F. & Wang, J. T. Characterization of integrative and conjugative element ICE*Kp1*-associated genomic heterogeneity in a *Klebsiella pneumoniae* strain isolated from a primary liver abscess. J. Bacteriol. 190, 515–526 (2008).

10. Lam, M. C. C. et al. Tracking key virulence loci encoding aerobactin and salmochelin siderophore synthesis in *Klebsiella pneumoniae*. Genome Med. 10, 77 (2018).

11. Lam, M. M. C. et al. Population genomics of hypervirulent *Klebsiella pneumoniae* clonal-group 23 reveals early emergence and rapid global dissemination. *Nat*. Comms 9, (2018).

12. Nassif, X., Fournier, J., Arondel, J. & Sansonetti, P. J. Mucoid Phenotype of *Klebsiella pneumoniae* Is a Plasmid-Encoded Virulence Factor. Infect Immun 57, 546–552 (1989).

13. Hsu, C. R., Lin, T. L., Chen, Y. C., Chou, H. C. & Wang, J. T. The role of *Klebsiella pneumoniae rmpA* in capsular polysaccharide synthesis and virulence revisited. Microbiology 157, 3446–3457 (2011).

14. Cheng, H. Y. et al. RmpA regulation of capsular polysaccharide biosynthesis in *Klebsiella pneumoniae* CG43. J Bacteriol. 192, 3144–3158 (2010).

15. Walker, K. A. et al. A *Klebsiella pneumoniae* Regulatory Mutant Has Reduced Capsule Expression but Retains Hypermucoviscosity. MBio 10, e00089–19 (2019).

16. Walker, K. A., Treat, L. P., Sepúlveda, V. E. & Miller, V. L. The Small Protein RmpD Drives Hypermucoviscosity in *Klebsiella pneumoniae*. MBio 11, e01750–20 (2020).

17. Ovchinnikova, O. G. et al. Hypermucoviscosity Regulator RmpD Interacts with Wzc and Controls Capsular Polysaccharide Chain Length. MBio 14, e00800–23 (2023).

18. Whitfield, C., Wear, S. S. & Sande, C. Assembly of Bacterial Capsular Polysaccharides and Exopolysaccharides. Annu. Rev. Microbiol. 74, 521–543 (2020).

19. Russo, T. A. et al. Identification of biomarkers for the differentiation of hypervirulent *Klebsiella pneumoniae* from classical *K. pneumoniae*. J Clin Microbiol **Online**, (2018).

20. Lam, M. M. C. et al. A genomic surveillance framework and genotyping tool for *Klebsiella pneumoniae* and its related species complex. Nat. Commun. 12, 4188 (2021).

21. Jenney, A. W. et al. Seroepidemiology of *Klebsiella pneumoniae* in an Australian Tertiary Hospital and Its Implications for Vaccine Development. J. Clin. Microbiol. 44, 102–107 (2006).

22. Sands, K. et al. Characterization of antimicrobial-resistant Gram-negative bacteria that cause neonatal sepsis in seven low- and middle-income countries. Nat. Microbiol. 6, 512–523 (2021).

23. Wyres, K. L. et al. Distinct evolutionary dynamics of horizontal gene transfer in drug resistant and virulent clones of *Klebsiella pneumoniae*. PLoS Genet. 15, e1008114 (2019).

24. Stamatakis, A. RAxML-VI-HPC: Maximum likelihood-based phylogenetic analyses with thousands of taxa and mixed models. Bioinformatics 22, 2688–2690 (2006).

25. Argimón, S. et al. Rapid Genomic Characterization and Global Surveillance of *Klebsiella* Using Pathogenwatch. Clin. Infect. Dis. 73, S325–S335 (2021).

26. Wick, R. R., Schultz, M. B., Zobel, J. & Holt, K. E. Bandage: interactive visualization of de novo genome assemblies. Bioinformatics 31, 3350–3352 (2015).

27. Seemann, T. Prokka: Rapid prokaryotic genome annotation. Bioinformatics 30, 2068–2069 (2014).

28. Gilchrist, C. L. M. & Chooi, Y.-H. clinker & clustermap.js: automatic generation of gene cluster comparison figures. Bioinformatics 37, 2473–2475 (2021).

29. Obrist, M. W. & Miller, V. L. Low copy expression vectors for use in *Yersinia* sp. and related organisms. Plasmid 68, 33–42 (2012).

30. Michelle, P., A., B. C., A., W. K. & L., M. V. A Serendipitous Mutation Reveals the Severe Virulence Defect of a *Klebsiella pneumoniae fepB* Mutant. mSphere 2, 10.1128/msphere.00341-17 (2017).

31. A., B. M., et al. Genome-Wide Identification of *Klebsiella pneumoniae* Fitness Genes during Lung Infection. MBio 6, 10.1128/mbio.00775-15 (2015).

32. Lawlor, M. S., Hsu, J., Rick, P. D. & Miller, V. L. Identification of *Klebsiella pneumoniae* virulence determinants using an intranasal infection model. Mol. Microbiol. 58, 1054–1073 (2005).

33. Brisse, S. et al. Virulent clones of *Klebsiella pneumoniae*: Identification and evolutionary scenario based on genomic and phenotypic characterization. PLoS One 4, (2009).

34. Liao, W., Liu, Y. & Zhang, W. Virulence evolution, molecular mechanisms of resistance and prevalence of ST11 carbapenem-resistant *Klebsiella pneumoniae* in China: A review over the last 10 years. J. Glob. Antimicrob. Resist. 23, 174–180 (2020).

35. Yi-Chyi, L., Hwei-Ling, P. & Hwan-You, C. RmpA2, an Activator of Capsule Biosynthesis in *Klebsiella pneumoniae* CG43, Regulates K2 *cps* Gene Expression at the Transcriptional Level. J. Bacteriol. 185, 788–800 (2003).

36. Lam, M. M. C. et al. Genetic diversity, mobilisation and spread of the yersiniabactin-encoding mobile element ICE*Kp* in *Klebsiella pneumoniae* populations. Microb Genom **Jul** 9, (2018).

37. Le, M. N.-T. et al. Genomic epidemiology and temperature dependency of hypermucoviscous *Klebsiella pneumoniae* in Japan. *Microb*. Genomics 8, (2022).

