## Additional File 2 for "Genomic and functional analysis of *rmp* locus variants in *Klebsiella pneumoniae*"

**Additional File 2.** Primers used in this work

**Name	Sequence^a^(5’→3’)	Use^b^**

KW531	GGCCCCCCCTCGAGGTCGACGATGAATATTGATGGATCAAAG	F pKW215, pKW222

KW532	GCTCTAGAACTAGTGGATCCGGAAACAAAAAGCTATACCATC	R pKW215

KW533	GGCCCCCCCTCGAGGTCGACGATGAATATTGATGGAGCAAAG	F pKW216

KW534	GCTCTAGAACTAGTGGATCCGGAAACCAAAAGTTATACCATC	R pKW216, pKW222, pLPT059

KW543	CGGGCCCCCCCTCGAGGTCGACGATGAATATTGATGGTTCAAAG	F pLPT059

_____________________________________________________________________________________________

^a^ Blue nt are overhangs for Gibson cloning. Underlined nt indicate restriction sites.

^b^ F, forward primer; R, reverse primer
