## Additional File 3 for "Genomic and functional analysis of *rmp* locus variants in *Klebsiella pneumoniae*"

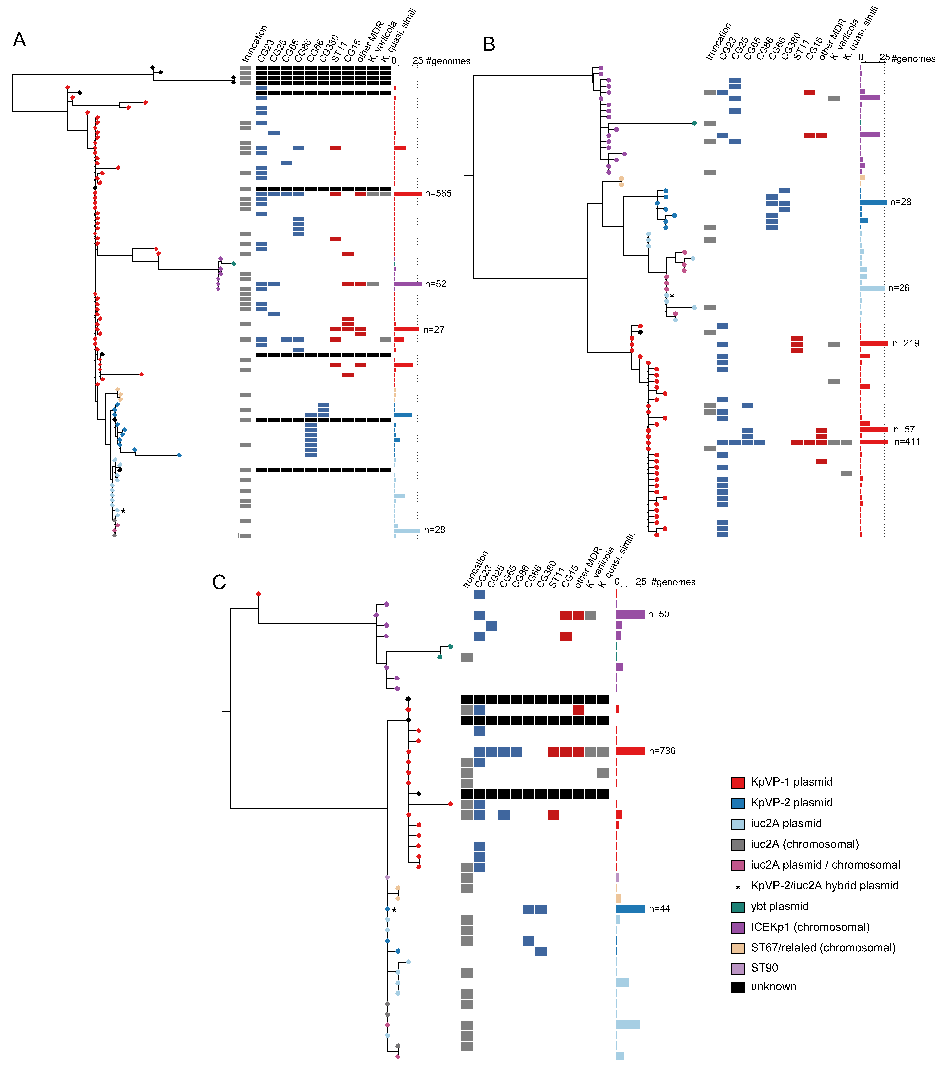

**Additional File 3. Phylogenetic relationships of the A. *rmpA,* B. *rmpD* and C. *rmpC* genes.** Each tree is a maximum-likelihood phylogeny with each node corresponding to an allelic variant. Nodes are coloured by the associated mobile genetic element according to the legend. Columns are as follows: presence or absence of a truncation, detection within a hypervirulent (blue) or MDR (red) clone or non-*K. pneumoniae* species. The number of genomes from which each allelic variant was detected is shown in the bar graph on the right-hand side.
