## Additional File 7 for "Genomic and functional analysis of *rmp* locus variants in *Klebsiella pneumoniae*"

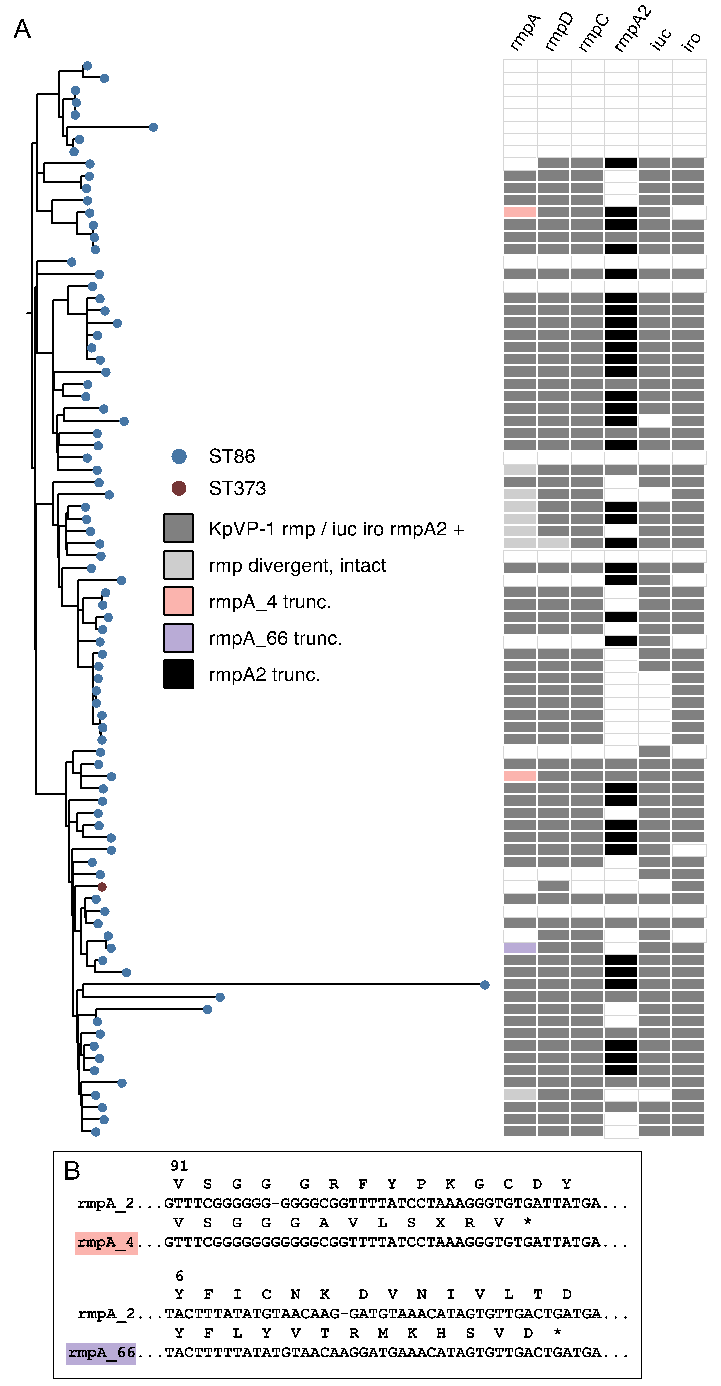

**Additional File 7. Distribution of *rmpADC* allelic variants and KpVP-1 associated virulence loci *rmpA2*, *iuc* and *iro* in *Klebsiella pneumoniae* clonal group 86. A.** Tree is a cgMLST SNP-based neighbour joining tree for 88 CG86 genomes generated from Pathogenwatch. Nodes are coloured by ST as per legend. Column shows the presence or absence (coloured white) of *rmpA, rmpD, rmpC, rmpA2, iuc* and *iro*, and allelic variant or truncation status as specified (coloured according to inset legend). **B.** Nucleotide and amino acid alignments are shown for two allelic variants with indels (i.e. *rmpA_4* and *rmpA_66*) compared to the wild-type intact variant (i.e. *rmpA_2*). Asterisk denotes premature stop codon.
